# Data augmentation based on dynamical systems for the classification of brain states

**DOI:** 10.1101/2020.01.08.898999

**Authors:** Yonatan Sanz Perl, Carla Pallavicini, Ignacio Perez Ipiña, Morten Kringelbach, Gustavo Deco, Helmut Laufs, Enzo Tagliazucchi

**Author notes:** Corresponding author Email address (YSP); (ET) (Enzo Tagliazucchi). These authors contributed equally to this work (Yonatan Sanz Perl, Carla Pallavicini).

## Abstract

The application of machine learning algorithms to neuroimaging data shows great promise for the classification of physiological and pathological brain states. However, classifiers trained on high dimensional data are prone to overfitting, especially for a low number of training samples. We describe the use of whole-brain computational models for data augmentation in brain state classification. Our low dimensional model is based on nonlinear oscillators coupled by the empirical SC of the brain. We use this model to enhance a dataset consisting of functional magnetic resonance imaging recordings acquired during all stages of the human wake-sleep cycle. After fitting the model to the average FC of each state, we show that the synthetic data generated by the model yields classification accuracies comparable to those obtained from the empirical data. We also show that models fitted to individual subjects generate surrogates with enough information to train classifiers that present significant transfer learning accuracy to the whole sample. Whole-brain computational modeling represents a useful tool to produce large synthetic datasets for data augmentation in the classification of certain brain states, with potential applications to computer-assisted diagnosis and prognosis of neuropsychiatric disorders.

## 1. Introduction

The discovery of non-invasive neuroimaging tools opened the way to the inference of the hidden brain states that are associated with observable behaviors. For this purpose, techniques such as functional magnetic resonance imaging (fMRI) provide high dimensional spatiotemporal data that can be used as the input for machine learning classifiers [1]. In these algorithms the parameters are learned from a training sample, and the resulting accuracy is then estimated from out-of-the-sample data. A sufficiently large number of examples is critical for successful training (i.e. avoiding overfitting), but the availability of neuroimaging data can be limited for certain rare neuropsychiatric conditions and for classifiers aimed at distinguishing between several groups of patients. While pooling data acquired in different laboratories can help alleviate this issue, it has been shown that heterogeneous experimental conditions can reduce the accuracy of the classifiers [2].

Data augmentation is a technique based on applying certain transformations to the available data with the objective of producing new surrogate training examples. In the case of image classification, for instance, these transformations may include rotations and shear mappings [3]. It is less obvious how to choose transformations that produce meaningful surrogate examples in the case of high dimensional spatiotemporal data, such as that provided by fMRI experiments. Faced with a similar problem, Tubaro and Mindlin recently proposed the use of low dimensional dynamical systems for data augmentation in deep learning [4]. Here, we have followed the analogous procedure of developing semi-empirical models of whole-brain activity for data augmentation in the classification of temporally extended brain states.

The computational models we developed and implemented receive as input several independent sources of empirical data, and can be optimized to reproduce observables derived from fMRI recordings [5]. A low dimensional dynamical system can be assigned to each region within a brain parcellation, and inter-regional coupling can be estimated from diffusion tensor imaging (DTI) data [6]. Using the normal mode of a Hopf bifurcation results in local dynamics with a transition from a fixed point towards a limit cycle, and in global dynamics coupled by the density of long-range white matter tracts [7, 8]. Finally, to reduce the dimension of the models, the local bifurcation parameters can be constrained by different functionally coherent brain systems, known as resting state networks (RSN) [9].

In the following, we show that these models reproduce the empirical correlation matrices between regional fMRI time series (also known as functional connectivity [FC] matrices), and that surrogate instances of FC matrices can be used for data augmentation in the problem of classifying the different stages of the human wake-sleep cycle. For this, synthetic time series were generated from the low dimensional models fitted to average and individual FC, which were used afterwards as input for multivariate random forest classifiers.

## 2. Material and Methods

### Participants and experimental protocol

63 healthy subjects participated in the original experiments (36 females, mean ± SD age of 23 ± 43.3 years). Written informed consent was obtained from all participants. The experimental protocol was approved by the local ethics committee (Goethe-Universität Frankfurt, Germany, protocol number: 305*/*07). The subjects were reimbursed for their participation. All experiments were conducted in accordance with the relevant guidelines and regulations, including the Declaration of Helsinki.

Participants entered the scanner in the evening (within half an hour of 7 PM) and underwent a resting state fMRI session with simultaneous EEG acquisition lasting for 52 minutes. Participants were not instructed to fall asleep, but were asked to relax, close their eyes and not actively fight the onset of sleep. Lights were dimmed in the scanner room and subjects were shielded from scanner noise using earplugs. The day of the study all participants reported a wake-up time between 5 : 00 AM and 11 : 00 AM, and a sleep onset time between 10 : 00 PM and 2 : 00 AM for the night prior to the experiment. Sleep diaries confirmed that these values were representative of the 6 days prior to the experiment.

### Simultaneous fMRI and EEG data collection

An optimized polysomnographic setting was employed to acquire electroencephalography (EEG) and electromyography (EMG) for sleep staging. Scalp potentials measured with EEG determine the classification of sleep into 4 stages (wakefulness, N1, N2 and N3 sleep) according to the rules of the American Academy of Sleep Medicine [10]. We selected a subset of 15 subjects who reached stage N3 sleep (i.e. deep sleep). Previous publications based on this dataset can be referenced for further details [11].

### Structural connectivity

Structural connectivity (SC) was obtained applying diffusion tensor imaging (DTI) to diffusion weighted imaging (DWI) recordings from 16 healthy right-handed participants (11 men and 5 women, mean age: 24.75 ± 2.54 years) recruited online at Aarhus University, Denmark. For each participant, a 90×90 SC matrix was obtained representing the density of white matter fiber tracts between regions of interest. The connectivity probability from a seed voxel *i* to another voxel *j* was defined as the proportion of fibers passing through voxel *i* that reached voxel *j* (sampling of 5000 streamlines per voxel) [12]. All the voxels in each region of the Automated Anatomical Labeling atlas (AAL [13]) were seeded (i.e. both grey and white matter voxels were considered). The connectivity probability *P*_*ij*_ from region *i* to region *j* was computed as the number of sampled fibers in region *i* that connected the two regions, divided by 5000 *n*, where *n* represents the number of voxels in region *i*. The resulting matrices were computed as the average across voxels within each region of interest in the AAL atlas, thresholded at 0.1 % (i.e. a minimum of five streamlines) and normalized by the number of voxels in the region. Finally, the data was averaged across all participants.

### Whole-brain models

We implemented a network of nonlinear oscillators coupled by the SC. Each oscillator was modeled using a normal form of a Hopf bifurcation and represents the dynamics at one of the 90 brain regions in the AAL atlas. In this type of bifurcation the qualitative nature of the solutions changes from a stable fixed point in phase space towards a limit cycle, allowing the model to represent the emergence of self-sustained oscillations. Thus, the key neurobiological assumption is that dynamics of macroscopic neural masses can range from fully synchronous to a stable asynchronous state governed by random fluctuations. We also assume that fMRI can capture the dynamics from both regimes with sufficient fidelity to be modeled by the equations.

Without coupling, the local dynamics of brain region *j* were modeled by the complex-valued equation:

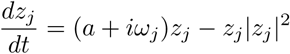

In this equation, *z* is a complex-valued variable (*z*_*j*_ = *x*_*j*_ + *iy*_*j*_), and *ω*_*j*_ is the intrinsic oscillation frequency of node *j*. The intrinsic frequencies ranged from 0.04 *-* 0.07 *Hz* and were determined by the averaged peak frequency of the bandpass-filtered fMRI signals of each individual brain region. The parameter *a* is known as the bifurcation parameter, and controls the dynamical behavior of the system. For *a* < 0 the phase space presents a unique stable fixed point at *z*_*j*_ = 0, thus the system decays asymptotically towards this point. For *a* > 0 the stable fixed point changes its stability, giving rise to a limit cycle and to self-sustained oscillations with frequency *f*_*j*_ = *ω*_*j*_*/*2*π* and amplitude proportional to the square root of *a* [7].

The coordinated dynamics of the resting state activity were modeled by a coupling term weighted by the SC. Nodes *i* and *j* were coupled by *C*_*ij*_ (the *i,j* entry of the SC matrix). To ensure oscillatory dynamics for *a* > 0, the SC matrix was scaled to a maximum of 0.2 (weak coupling assumption) [7]. In full form, the coupled differential equations of the model are the following:

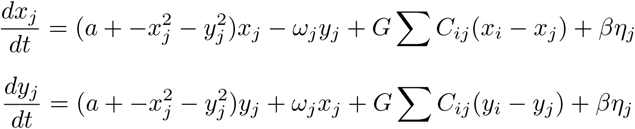

The parameter *G* represents a global factor that scales the SC equally for all the nodes. These equations were integrated to simulate empirical fMRI signals using the Euler-Maruyama algorithm with a time step of 0.1 seconds. *η*_*j*_ represents additive Gaussian noise in each node and was fixed at 0.04. When *a* is close to the bifurcation (*a* ≈ 0) the additive gaussian noise gives rise to complex dynamics as the system continuously switches between both sides of the bifurcation.

### Fitting the model to the empirical data

We used the group-averaged FC as the empirical observable to be fitted by the model. The fMRI signal from each region in the AAL atlas was filtered in the 0.04–0.07 *Hz* frequency range since, when mapped to the gray matter, this frequency band was shown to contain more reliable and functionally relevant information compared to other frequency bands, and to be less affected by physiological noise [14]. Subsequently, the filtered time series were transformed to z-scores. For each brain state, participants were selected based on the presence of uninterrupted epochs of that state lasting more than 200 samples, resulting in 15 participants. Afterwards, the FC matrix was defined as the matrix containing the correlation coefficients between the average fMRI signals from all pairs of regions of interest in the AAL atlas. Fixed-effect analysis was used to obtain group-level FC matrices, meaning that the Fisher’s R-to-z transform (*z* = *atanh*(*R*)) was applied to the correlation values before averaging across participants within each state of consciousness.

When applying the above described model to simulate the regional fMRI signals, we incorporated an anatomical prior based on 6 major RSN [9] with the objective of constraining how different groups of nodes could contribute to the local bifurcation parameters. In this way, we embedded the dynamics of the 90 independent regions into a 6-dimensional parameter space. Each bifurcation parameter was constructed as the linear combination of the 6 parameters associated with the RSN. Note that regions could belong to more than one RSN, and thus the bifurcation parameters could receive independent contributions from multiple RSN. We simulated 200 time samples for each subject, and then repeated the procedure described above to compute the simulated average FC matrices for each state. To simulate individual FC time series we increased the number of time samples to 2000. The goodness of fit was determined by the structure similarity index (SSIM) [15], an image similarity metric that factors both the similarity between the image means and between their covariance structures. The optimization procedure was based on genetic algorithms applied to infer the 6 parameters that maximize the goodness of fit. Further details can be found in previous work implementing the same model [16].

### Multivariate machine learning classifiers and data augmented by the model

We trained random forest classifiers [17] to distinguish sleep from wakefulness based on FC matrices, using a five-fold cross-validation procedure to estimate the accuracy. Classifiers were trained to distinguish between wakefulness and a certain sleep stage, and their accuracy was then tested in the classification between wakefulness and the same as well as other sleep stages (i.e. transfer learning accuracy).

Random forest classifiers were implemented using scikit-learn (https://scikit-learn.org/) [18]. Briefly, the random forest algorithm builds upon the concept of a decision tree classifier, where samples are iteratively split into two branches depending on the values of their features. For each feature, a threshold is introduced so that the samples are separated in a way that maximizes a metric of the homogeneity of the class labels assigned to each branch. The algorithm stops whenever a split results in a branch where all the samples belong to the same class, or when all features were already used for a split. Since this procedure is prone to overfitting, the random forest algorithm trains an ensemble of decision trees based on a randomly chosen subset of the features, and then computes the label prediction as the majority vote across all the individual trees.

We trained random forest classifiers with 1000 decision trees and a random subset of features of size equal to the (rounded) square root of the total number of features. The quality of each split in the decision trees was measured using Gini impurity, and the individual trees were expanded until all leaves were pure (i.e. no maximum depth). No minimum impurity decrease was enforced at each split, and no minimum number of samples was required at the leaf nodes of the decision trees (the classifier hyperparameters can be found in https://scikit-learn.org/).

To assess the statistical significance of the accuracy values, we trained and evaluated a total of 1000 random forest classifiers using the same features (i.e. FC matrices) but scrambling the class labels. We then constructed an empirical p-value by counting the how many times the accuracy of the classifier with scrambled class labels was greater than that of the original classifier. All accuracies were determined as the area under the receiver operating characteristic curve (AUC). Subsequently, the generalizability of the classifiers to distinguish other sleep states from wakefulness was evaluated by applying both the original and scrambled classifiers, and constructing a p-value in a similar way.

We repeated the aforementioned procedure using data augmentation, given by the output of the whole-brain computational model. The inclusion of additive noise in the model gives rise to different simulated time series for each independent run, and consequently to different FC matrices. We optimized model parameters using the average FC matrices as the targets, and used these parameters to simulate 100 surrogate samples for each instance of random forest classifier based on synthetic data.

In this way, we obtained 100 synthetic samples for each stage. Based on these surrogate samples, we trained classifiers to distinguish wakefulness from sleep and measured the accuracy of these classifiers using the empirical data. We also determined the transfer learning accuracy by evaluating the performance of the classifiers trained with surrogate data of a certain sleep stage in the problem of classifying empirical data corresponding to another sleep stage. Finally, we randomly selected three subjects from the empirical dataset and repeated this procedure using single subject FC matrices as optimization targets for the whole-brain model.

## 3. Results

The procedure we followed is outlined in Fig. 1. First, we combined three different sources of empirical data to inform the computational model based on coupled Hopf bifurcations, as explained in the Materials and Methods section. Then, we trained and evaluated random forest classifiers to distinguish different pairs of sleep stages based on empirical and synthetic fMRI data, as well as on single subject synthetic data.

**Figure 1:**
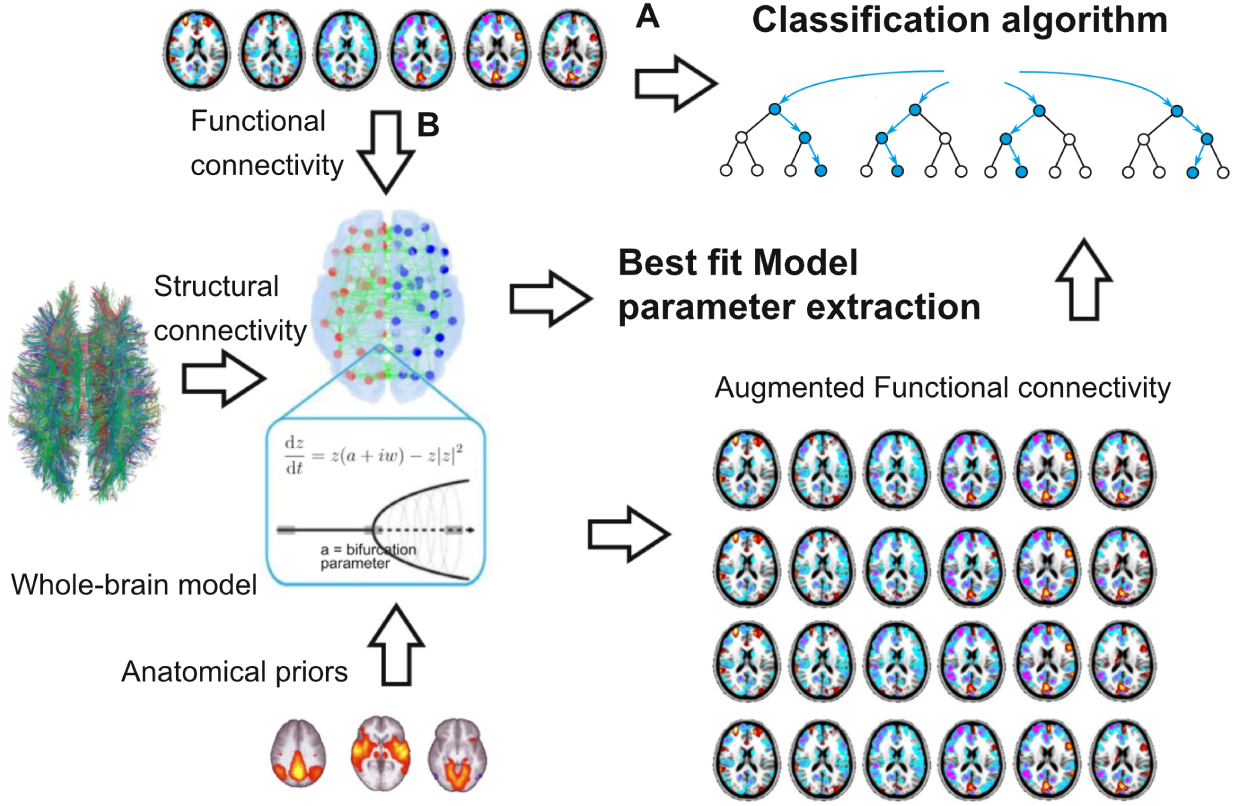
Outline of the procedure followed to generate the synthetic data and train the machine learning classifiers. Three sources of empirical data informed the computational model based on coupled nonlinear oscillators. SC represented the coupling strength between oscillators, FC was used as the target function for parameter optimization with genetic algorithms, and 6 RSN determined the anatomical priors constraining local contributions to bifurcation parameters. The empirical FC was used both as input to the random forest classifiers (A) and as target function (B) in the optimization procedure. After this step the model generated surrogate samples to augment the training data.

The first row of Fig. 2 shows the average empirical FC matrices corresponding to wakefulness and the three stages of NREM sleep (N1, N2, N3). In these matrices, rows and columns correspond to one of the 90 regions in the AAL atlas, and the correlation coefficient between fMRI time series is indicated by the color scale. The remaining rows show the FC matrices computed for three randomly chosen subjects.

**Figure 2:**
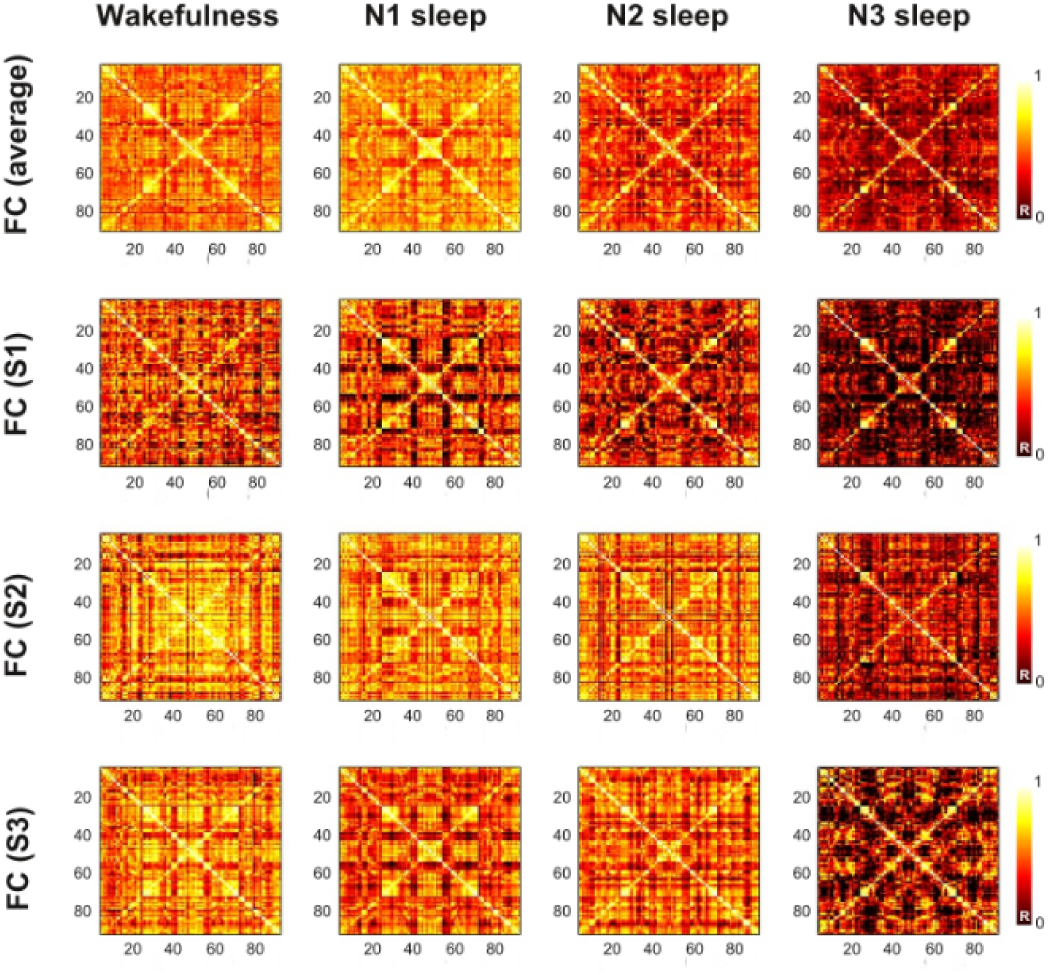
Empirical FC matrices containing the correlation coefficients (R) between fMRI time series from all pairs of regions in the AAL atlas. The first row displays the average FC matrices for wakefulness, N1, N2 and N3 sleep. The other rows contain the same information for three randomly chosen individuals.

Fig. 3 contains the same information computed from the synthetic fMRI data. The main difference between the empirical and simulated matrices appeared in the contradiagonal, which corresponds to interhemispheric (or homotopic) connections (i.e. connections between two regions symmetrically located with respect to the midline), a difference consistent with the observation that DTI tends to underestimate long-range fiber tracts [19]. In general, the model fit was better for the average FC compared to individual subjects (see Table 1).

**Table 1:** Goodness of fit (1-SSIM) between empirical and simulated FC matrices, both for the average data and for the three individual subjects (S1, S2, S3).

|  | **W** | **N1** | **N2** | **N3** |
| --- | --- | --- | --- | --- |
| Average | 0.60 | 0.60 | 0.59 | 0.63 |
| S1 | 0.73 | 0.72 | 0.76 | 0.79 |
| S2 | 0.63 | 0.70 | 0.46 | 0.78 |
| S3 | 0.58 | 0.70 | 0.61 | 0.73 |

**Figure 3:**
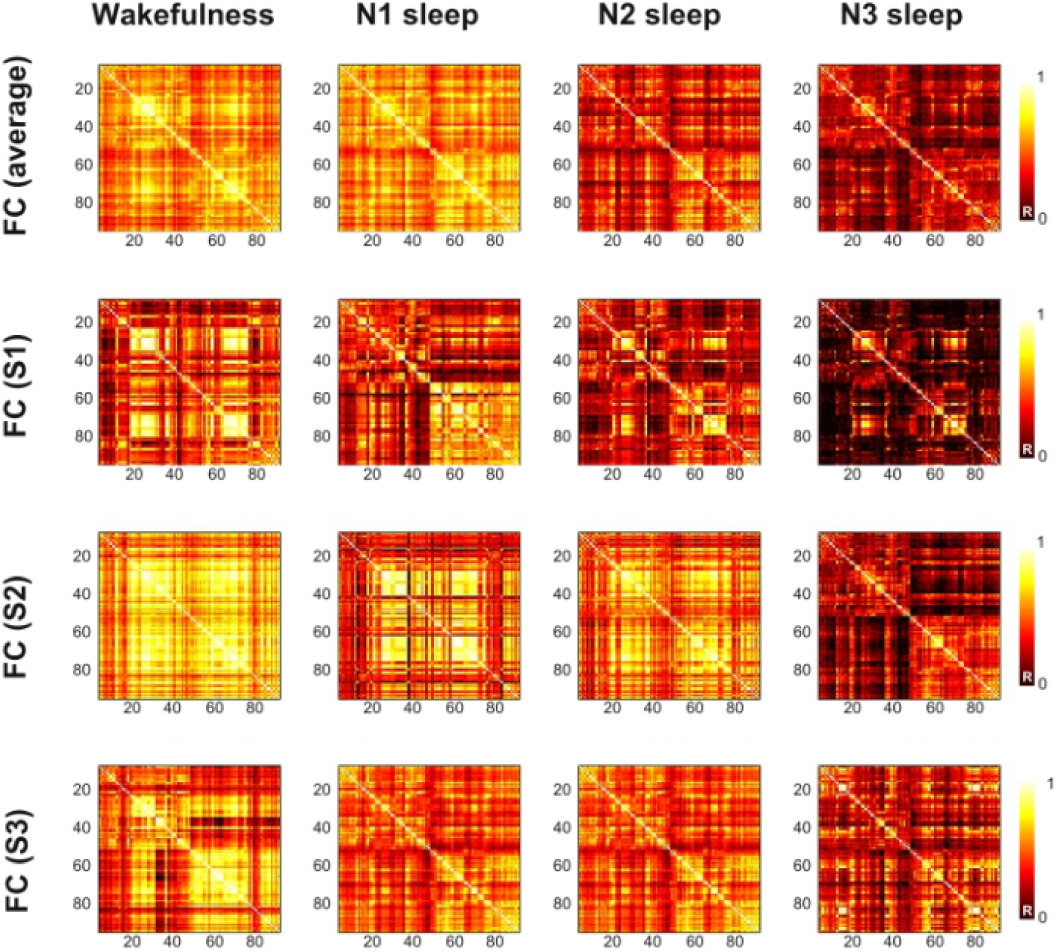
FC matrices containing the correlation coefficients (R) between synthetic fMRI time series from all pairs of regions in the AAL atlas. The first row displays the optimal simulated FC matrices for wakefulness, N1, N2 and N3 sleep. The other rows contain the same information for three randomly chosen individuals.

Panel A of Fig. 4 shows the histograms of AUC values representing the transfer learning accuracy for random forest classifiers trained using 100 synthetic wakefulness samples and 100 synthetic N1/N2/N3 sleep samples, and evaluated in the empirical data. Each column and row indicates the sleep stage used for training and testing, respectively. For instance, the second plot of the first row contains the AUC histograms obtained in the classification between the empirical FC matrices from wakefulness and N2 sleep (*N* = 15 subjects), using the classifier trained to distinguish wakefulness from N1 sleep based on synthetic fMRI data (*N* = 100 surrogates). The histograms in red correspond to the AUC values obtained using the real data, while the histograms in blue indicate the AUC values obtained after shuffling the data labels. Label shuffled is used as a null model to obtain the p-values shown in the insets.

**Figure 4:**
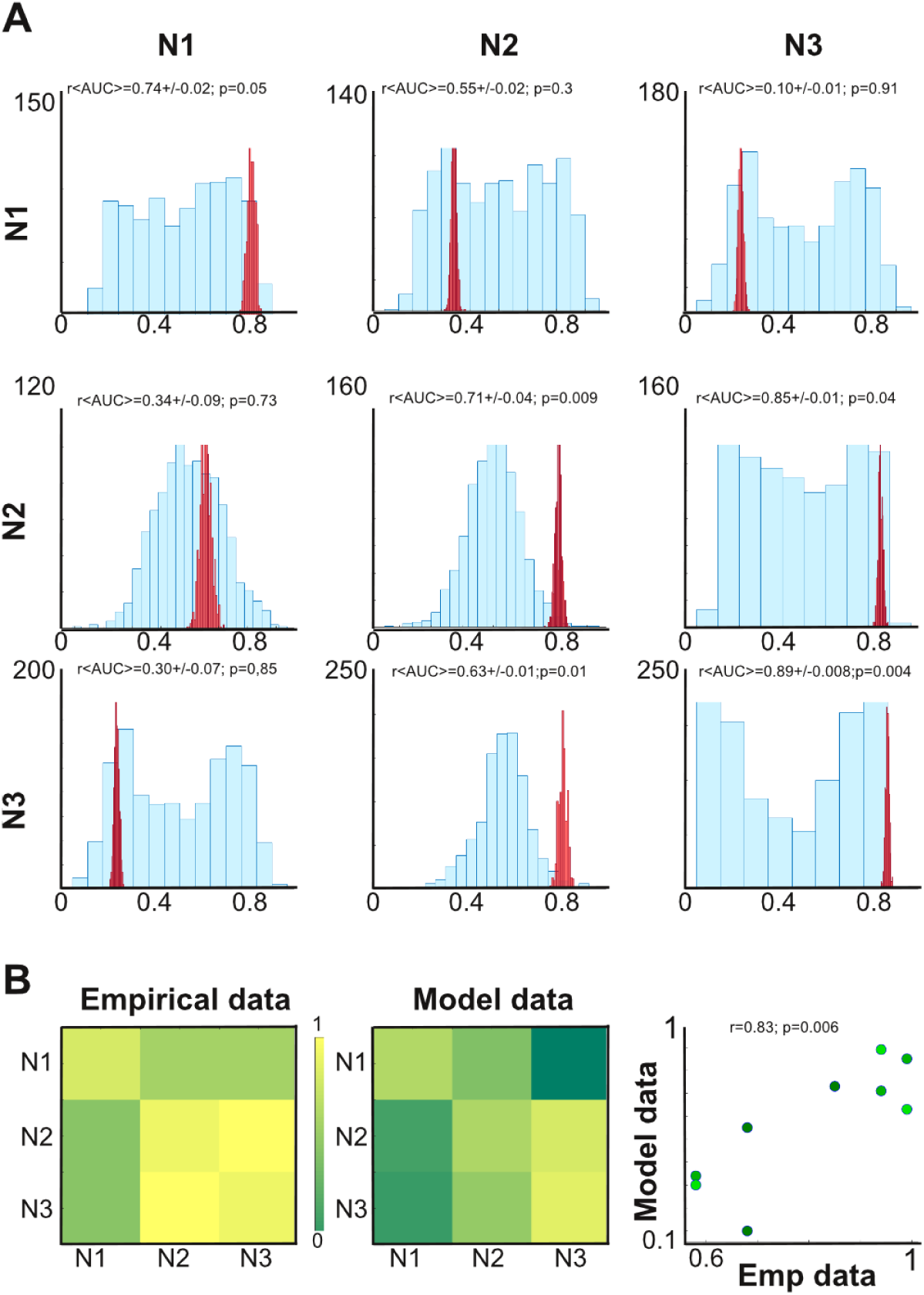
A) Histograms of AUC values for the random forest classifiers trained to distinguish wakefulness vs. the sleep stages indicated in the rows, and tested in the classification of wakefulness vs. the sleep stages indicated in the columns. All classifiers were trained using synthetic FC matrices fitted to the average FC matrices (*N* = 100 surrogates) and evaluated using the empirical data (*N* = 15). B) Matrices containing the average AUC values obtained for the random forest classifiers trained using the empirical (left) and the synthetic (right) data. The scatter plot contains the entries of the “empirical data” vs. the “model data” matrices.

The matrices in Panel B of Fig. 4 summarize the average AUC obtained for all training-generalization pairs. It is clear from observing these matrices that machine learning classifiers presented the highest transfer learning accuracy when generalizing between N2 and N3 sleep. This result was obtained using both synthetic (left) and empirical (right) data for training. The scatter plot compares the entries of both matrices, showing a positive correlation which supports the similarity between the empirical and simulated AUC matrices.

Fig. 5 shows that data augmentation based on single subject FC matrices can be used to train machine learning classifiers that present significant accuracy in the classification of wakefulness from N2 and N3 sleep. The rows correspond to random forest classifiers trained using data generated by computational models fitted to the empirical FC of S1, S2 and S3 (*N* = 100 surrogates). The columns indicate the sleep stage to be distinguish from wakefulness. All histograms contain AUC values obtained from the evaluation of these models on the empirical data (*N* = 15 subjects), both with unshuffled (red) and shuffled (blue) class labels. The resulting p-values indicate that classifiers trained using synthetic data from individual subjects can successfully generalize to the whole sample in the classification of wakefulness vs. N2 and N3 sleep, but not vs. N1 sleep.

**Figure 5:**
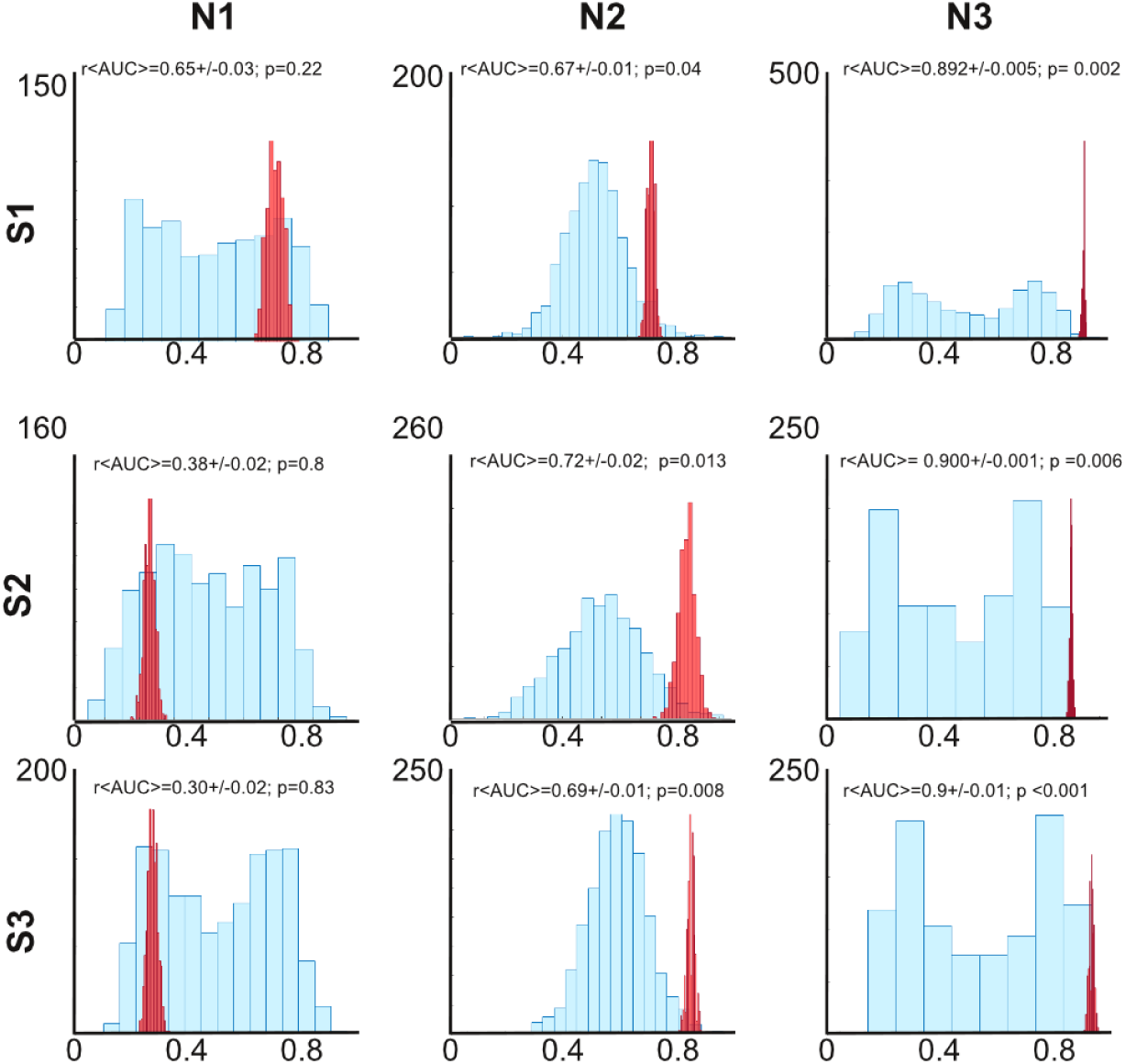
Histograms of AUC values for random forests classifiers trained to distinguish N1, N2 and N3 from wakefulness. Classifiers were trained using synthetic FC matrices based on data from individual subjects (*N* = 100 surrogates) and evaluated on empirical samples (*N* = 15 subjects). Histograms in red correspond to data without label shuffling, while blue indicates AUC after label shuffling. Insets contain the mean AUC ± SD and the associated p-values.

## 4. Discussion

One of the main limitations for the training of machine learning classifiers is the amount of available data. Training is generally successful provided sufficient data and informative features, but overfitting can drastically reduce generalization performance if only few training samples are available. Data augmentation techniques can attenuate this problem by introducing certain transformations (such as shear mappings and rotations, in the case of images); however, it is not clear how complex spatiotemporal data should be transformed to create meaningful surrogate samples. Following recently published work [4], we showed that low dimensional dynamical systems fitted to empirical observables can be successfully applied for data augmentation with the purpose of brain state classification. We note that surrogate time series can be produced by methods not based on dynamical systems (e.g. [20]). However, models such as the one we employed can represent advantages in terms of interpretation and conceptual clarity. They can also be tailored to train classifiers using synthetic samples deviating in useful ways from the available experimental data. Finally, their semi-empirical nature facilitates the transition towards the single subject level. In the following, we discuss these advantages in the context of the present work.

The computational models we used to generate synthetic data did not strive for biological realism, instead, we decided to focus on the simplest dynamics that could present the kind of behavior needed to provide the classifiers with representative surrogates for training. Based on this data, the classifiers were capable of inferring the optimal separation boundary between classes and presented significant transfer accuracy for the generalization between N2 and N3, which was expected considering the high behavioral and physiological similarities between these stages compared to N1 sleep [10]. The classifier transfer learning accuracy matrices obtained from synthetic and empirical data were very similar, supporting the conceptual validity of our simple model for whole-brain dynamics, which could be expected from previous work based on similar dynamics [7, 8, 16]. Low complexity models can simultaneously preserve the informativeness of the surrogates while allowing the exploration of a small set of interpretable parameters.

The classification of brain states based on fMRI recordings is a promising tool for the automated diagnosis and prognosis of certain neuropsychiatric patients [21], however, this promise is frequently undermined by small sample sizes [22]. Building databases of fMRI recordings can be costly and time consuming for diseases that are rare or difficult to investigate with neuroimaging. Also, developing algorithms for differential diagnosis requires multilabel classifiers, which further reduces the number of samples per class. A possible solution to this issue is gathering data from multiple research groups; however, different scanners and imaging sequences can be critical confounds for machine learning classifiers [2]. As an alternative, we proposed that adequate data augmentation techniques based on computational modeling can contribute to overcoming these limitations. We note that these are not mutually exclusive solutions, for instance, models could be use to explore parametrically how classifiers are confounded by factors related to variability in the experimental conditions.

The outcome of our model depends upon a relatively low number of parameters, which could be explored to train classifiers with surrogate samples including perturbations that represent the hypothesized outcome of certain interventions. For instance, the outcome of surgical brain resection in certain forms of epilepsy could be modeled by localized SC changes [23]. By artificially inducing these changes in the model parameters (including the structural coupling between nodes) it could be possible to produce synthetic data useful to train classifiers that can be applied to estimate the likelihood of success after the intervention. The same logic could be applied to other kinds of treatments, such as pharmacological interventions and non-invasive brain stimulation protocols, as well as to train machine learning classifier with data that simulates specific lesions, such as those arising from stroke and traumatic brain injury.

We have shown that data augmentation using models fitted to single subject FC matrices also allowed the classifiers to distinguish between wakefulness and sleep. As such, our results represent an encouraging proof of concept, but care should be exercise when attempting to generalize this result to other brain states. Since sleep is a physiological process and our population consisted of healthy participants, we expected that individual subjects could provide enough information to develop classifiers accurate at the group level. However, this cannot be taken for granted in surrogates obtained from models fitted to individual patients, where higher inter-subject variability may arise from abnormalities in brain structure and function. Since these limitations could be informative of such abnormalities, low dimensional whole-brain models should be further explored in the context of reproducing single subject FC from the individual SC of the patients [24].

In conclusion, we have shown that dynamical systems constitute a valuable tool for generating synthetic spatiotemporal data based on a small number of examples, a tool that can be naturally applied for data augmentation when training automated classifiers using fMRI data. Future work should study the possibility of overcoming data scarcity in other systems that can be modeled by simple dynamics, contributing to the fruitful cross-fertilization of artificial intelligence and physics.

